# PH-SENSITIVE NANODROPLETS FOR CONTROLLED DELIVERY OF BERBERINE CHLORIDE

**DOI:** 10.1101/2024.03.18.585497

**Authors:** Shengjie Hao, Shiyu Wang, Jiaqi Cao, Zhiyuan Xue, Zeyu Luo, Peirun Wu, Guilin Chen

## Abstract

**Background:** The development of nanocarriers with precise control over drug release is crucial for targeted therapy. This study focuses on the design and optimization of pH-sensitive gelatin/perfluorohexane (PFH) nanodroplets loaded with berberine chloride, a model drug relevant to traditional Chinese medicine.

**Subjects and Methods:** Nanodroplets were prepared using an emulsion technique, with optimization of parameters including homogenization rate, polymer concentration, surfactant, drug, and perfluorocarbon conte nt.

**Results:** The optimized formulation resulted in nanodroplets with a mean particle size of 281.7 nm and a drug encapsulation efficiency of 66.8 ± 1.7%. Characterization studies confirmed successful encapsulation and pH-responsive behavior. Ultrasound stimulation significantly enhanced drug release, with 150 kHz frequency proving more effective than 1 MHz. Stability studies demonstrated prolonged stability over one month at 4°C. Following 10 minutes of ultrasound irradiation, the nanodroplets exhibited 89.4% cumulative drug release.

**Conclusions:** In conclusion, these pH-sensitive nanodroplets show potential for delivering berberine chloride in a controlled manner, connecting traditional Chinese medicine with contemporary drug delivery methods.

## 1. Introduction

Recent advancements in nanotechnology have ushered in a new era in drug delivery systems, prominently featuring the utilization of nanocarriers tailored for precise and controlled drug release. Nanocarriers, which represent nano-sized constructs meticulously engineered to encapsulate and ferry therapeutic agents such as drugs and genes, offer unparalleled flexibility with their composition of myriad materials including lipids, polymers, metals, or hybrid formulations. [1],[2]. Their diminutive size and augmented surface-to-volume ratio confer significant advantages in drug delivery, facilitating enhanced drug penetration, heightened cellular uptake, and precise interaction with target cells and tissues, ultimately leading to more efficacious therapeutic out comes [3]. Through the synergistic application of ultrasound energy and responsive materials, drug delivery has under gone a revolution, enabling precise targeting with reduced side effects and heightened treatment efficacy. This paradigm allows for the controlled release of encapsulated therapeutic agents, precisely localized to target tissues through external stimuli. Ultrasound not only facilitates precise targeting but also enhances drug penetration via sonoporation, thereby fostering enhanced cellular and tissue interaction. The amalgamation of ultrasound and nanotechnology offers a non-invasive and patient-friendly approach, enhancing the quality of life for patients undergoing treatment and paving the way for personalized medicine[4].

Liquid perfluorocarbon (PFC) nanoemulsions, known for their chemical inertness, biocompatibility, and superior oxygen and gas-carrying capabilities, serve as promising carriers for therapeutic agents. When formulated into nanoemulsions, PFCs exhibit unique properties that can be triggered by external stimuli. Ongoing research in PFC nanoemulsions focuses on optimizing their formulation, exploring novel stimuli-responsive behaviors, and devising strategies for targeted drug delivery to augment therapy efficacy and mitigate adverse effects[5].Low-boiling point perfluorocarbon nanodroplets, such as perfluorohexane (PFH) droplets, have garnered significant interest in biomedical applications, particularly in ultrasound imaging and therapy[6]. The low-boiling point characteristic of PFH nanodroplets denotes their propensity to transition from liquid to gas phase at relatively low temperatures compared to other substances, enabling rapid phase transitions upon exposure to specific stimuli, such as ultrasound. When subjected to ultrasound, PFH nanodroplets undergo acoustic droplet vaporization (ADV), swiftly vaporizing to form gas-filled microbubbles in response to ultrasound waves[7].The utilization of PF H nanodroplets stimulated by ultrasound presents a promising avenue for cancer drug delivery. These nanodroplets can be engineered to accumulate specifically at tumor sites, where they undergo vaporization upon ultrasound triggering, facilitating localized drug release within the tumor microenvironment. This targeted drug delivery approach minimizes systemic exposure and diminishes off-target effects, thereby optimizing therapeutic outcomes[8]. Moreover, ultrasound affords precise control over the timing and extent of drug release, enhancing therapeutic efficacy while mitigating toxicity concerns. Additionally, the vaporization of PFH nanodroplets induces mechanical effects that augment drug penetration into tumor tissues, amplifying the effectiveness of chemotherapy. The synergistic combination of PFH nanodroplets and ultrasound with other therapeutic modalities offers a non-invasive, repeatable approach that aligns with existing ultrasound technology, promising enhanced treatment outcomes for cancer patients[9] .However, while PFH nanodroplets show considerable potential for drug delivery applications, several drawbacks merit consideration. Firstly, their limited payload capacity poses challenges for delivering high doses or large therapeutic molecules. Secondly, their short circulation half-life necessitates repeated dosing or alternative administration strategies to maintain therapeutic concentrations[10]. Additionally, concerns regarding immunogenicity and stability must be addressed to ensure their safety and efficacy. Despite these challenges, strategies such as incorporating chitosan or alginate shells have shown promise in enhancing stability and enabling targeted drug delivery. Chitosan, in particular, offers electrostatic interactions that stabilize nanodroplets and facilitate controlled drug release, while alginate provides pH-sensitive properties for targeted release in the acidic tumor microenvironment. However, the moderate stability of chitosan and alginate may impact long-term storage and shelf life, necessitating careful consideration during formulation design [11].Gelatin emerges as a compelling alternative for PFH nanodroplet shells, offering excellent stability, mechanical properties, and biocompatibility. Its versatility allows for easy modification to enhance specific properties such as drug loading capacity, controlled release, and targeting abilities[12]. Furthermore, gelatin’s compatibility with various fabrication techniques enables the efficient production of uniform nanodroplets, with its biodegradability and non-toxic nature supporting its suitability for biomedical applications.

In conclusion, gelatin exhibits substantial potential as a shell material for PFH nanodroplets, facilitating their successful deployment in drug delivery and other therapeutic endeavors [13].In this study, we developed ultrasound-responsive nanodroplets as an innovative smart drug delivery system tailored for nano-therapy, particularly for diseases like cancer. Our focus was on fabricating multifunctional smart gelatin/perfluorohexane nanodroplets loaded with berberine chloride for targeted drug delivery. Berberine chloride, an alkaloid naturally extracted from plants such as Berberis vulgaris and Coptis chinensis, holds a long-standing tradition in Chinese medicine for its antibacterial, antifungal, and anti-inflammatory properties. Recently, its potential as an anti-cancer agent has garnered increasing attention due to its efficacy in inhibiting cancer cell proliferation, inducing apoptosis, and sensitizing cancer cells to chemotherapy by targeting multiple signaling pathways implicated in cancer development. Concerns over poor solubility and low bioavailability have hindered its clinical utility [14].The encapsulation of berberine chloride within gelatin/perfluorohexane nanodroplets addresses these limitations by enhancing its solubility and stability, thereby improving drug dispersion and preventing precipitation. Gelatin forms a protective barrier around berberine chloride, shielding it from degradation and enzymatic breakdown in biological systems, ensuring sustained release and optimal therapeutic effect. The diminutive size and surface properties of gelatin/perfluorohexane nanodroplets promote accumulation at the target site via the enhanced permeability and retention (EPR) effect, facilitating preferential accumulation in tumor tissues for enhanced therapeutic activity.

## 2. Materials and Methods

### 2.1 Materials

Berberine chloride and perfluorohexane, supplied by Lanzhou Veterinary Institute (Lanzhou, China), were used in the experiments. Gelatin (type B, Bloom 250, G108395), Tween 80 (T104866) as a surfactant, and glutaraldehyde (2 5% in H2O, G105906) were obtained from Aladdia (Shanghai, China). Additionally, sodium hydroxide was purchased from the same supplier. Deionized water was utilized throughout all procedures, while all other chemicals used w ere of reagent grade.

### 2.2 Preparation of PFH Nanodroplets with a Gelatin Shell Containing Berberine Chloride

Nanodroplets were synthesized using the nano-emulsion method, with modifications to our previous methodology to suit the current study 11. Initially, gelatin solutions of var ying concentrations (Table 1) were prepared in deionized water by stirring on a magnetic stirrer (Biometer) at 50°C and 500 rpm for 10 minutes until complete clarity was achieved. The pH was adjusted to 6 with 0.1 M NaOH to prevent gelatin deposition. After cooling to room temperature, the polymeric solutions underwent filtration. Subsequently, nanodroplets were synthesized using the nano-emulsion method with modifications to the previous methodology for the current study. Berberine chloride was dissolved in deionized water using a magnetic stirrer, and a surfact ant solution (0.5% v/v Tween 80) was added to perfluoro hexane, followed by the drug solution. This mixture was homogenized for 2 minutes at 24,000 rpm using an Ultra

**Table 1.**
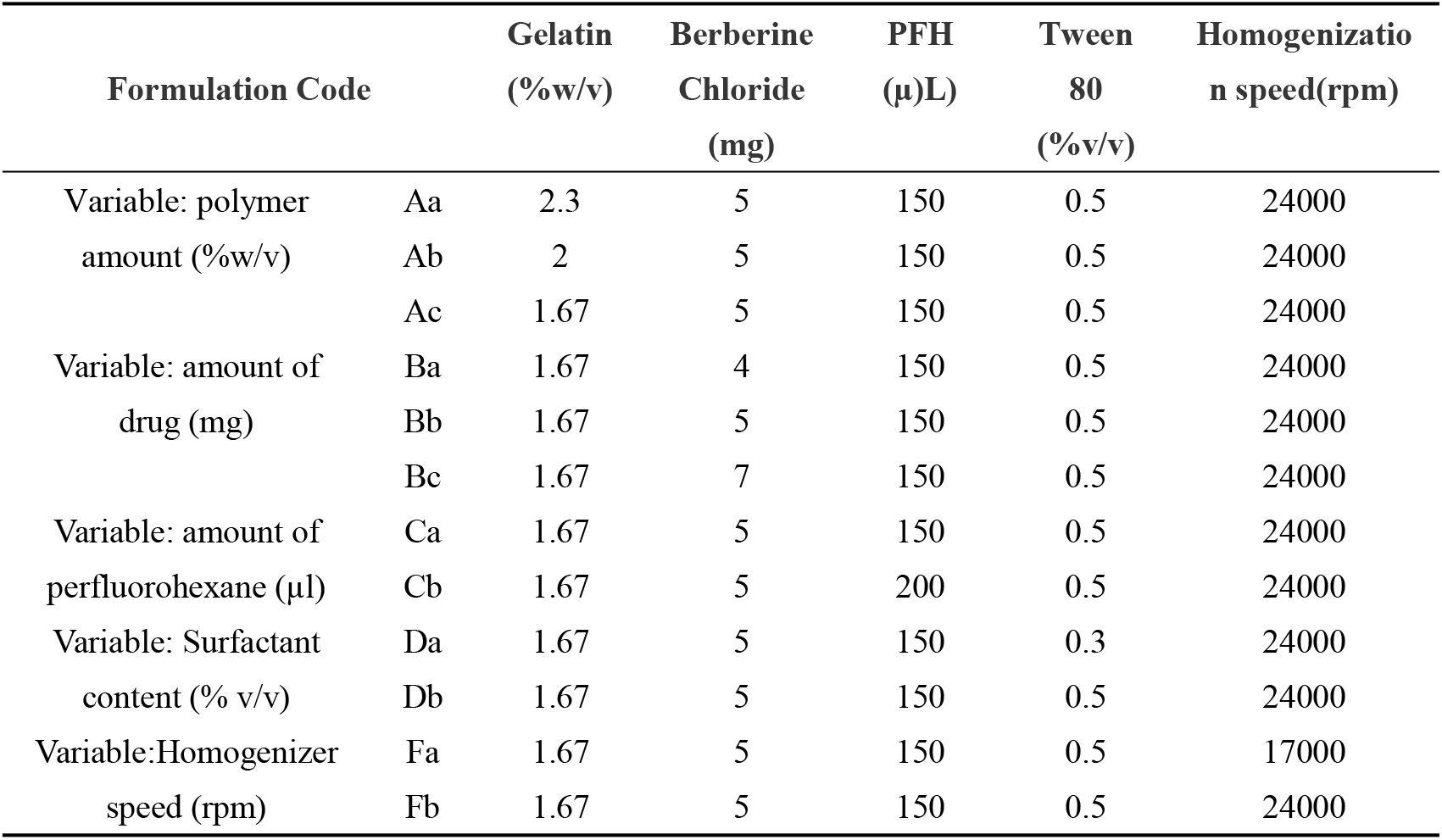
Formulations and process variables on the production of nanodroplets loaded with berberine chloride using PFH and gelatin. (Add 3 milliliters of deionized water to all components)

| Formulation Code | | Gelatin<br>(%w/v) | Berberine<br>Chloride<br>(mg) | PFH<br>(μL) | Tween<br>80<br>(%v/v) | Homogenization<br>speed(rpm) |
| --- | --- | --- | --- | --- | --- | --- |
| Variable: polymer<br>amount (%w/v) | Aa | 2.3 | 5 | 150 | 0.5 | 24000 |
|  | Ab | 2 | 5 | 150 | 0.5 | 24000 |
|  | Ac | 1.67 | 5 | 150 | 0.5 | 24000 |
| Variable: amount of<br>drug (mg) | Ba | 1.67 | 4 | 150 | 0.5 | 24000 |
|  | Bb | 1.67 | 5 | 150 | 0.5 | 24000 |
|  | Bc | 1.67 | 7 | 150 | 0.5 | 24000 |
| Variable: amount of<br>perfluorohexane (μl) | Ca | 1.67 | 5 | 150 | 0.5 | 24000 |
|  | Cb | 1.67 | 5 | 200 | 0.5 | 24000 |
| Variable: Surfactant<br>content (% v/v) | Da | 1.67 | 5 | 150 | 0.3 | 24000 |
|  | Db | 1.67 | 5 | 150 | 0.5 | 24000 |
| Variable:Homogenizer<br>speed (rpm) | Fa | 1.67 | 5 | 150 | 0.5 | 17000 |
|  | Fb | 1.67 | 5 | 150 | 0.5 | 24000 |

### 2.3 Characterization of Berberine Chloride-Loaded Nanodroplets

#### 2.3.1 Particle Size, Size Distribution and Zeta Potential

The average particle size, polydispersity index (PDI), and zeta potential of the nanodroplets were determined using dynamic light scattering (DLS) with a zetasizer nano zs90 (Malvern, Shanghai). Prior to measurement, the nanodroplet emulsions were diluted 1:4 with deionized water. Subsequently, particle size, size distribution, and zeta potential were analyzed to gain insights into the properties and behavior of the Berberine Chloride-loaded nanodroplets. This characterization step provides crucial information for understanding the stability and efficacy of the formulation, shedding light on its potential applications in pharmaceutical and biomedical fields.

#### 2.3.2 Morphology of Drug-Loaded Nanodroplets

Transmission electron microscopy (TEM) was utilized to visualize the morphology and structure of the nanodroplets. In sample preparation, a drop of the emulsion was deposited onto a 400-mesh carbon-coated copper grid and air-dried at room temperature. Subsequently, to improve contrast for imaging, the samples were treated with 1% alkaline phosphotungstic acid (PTA) for several minutes before ana lysis.

#### 2.3.4 Fourier Transform Infrared (FTIR) Spectroscopy

FTIR spectroscopy was used to analyze the chemical com position and identify specific functional groups of the drug delivery system. Nanodroplets were lyophilized to produce micronized KBr pellets, and spectra were recorded in the 4500–400 cm−1 range.

#### 2.3.4 Evaluation of Encapsulation Efficiency

Turrax T25 homogenizer. The polymer solution was then added dropwise to the emulsion under homogenization at 13,000 rpm for 5 minutes. Glutaraldehyde solution, a crosslinking agent, was added dropwise and stirred at 5000 rpm. The crosslinking process was stopped by adding 8 mL of aqueous sodium metabisulfite solution (1.6% w/v). To obtain nanodroplets with the desired size in the nanorange and high drug loading efficiency, various formulations were tested by adjusting different process parameters. Nanodroplet size, drug loading efficiency, and drug release kinetics were influenced by these parameters, including polymer concentration, drug content, surfactant concentration, and perfluorohexane amount. Additionally, the study explored process parameters such as homogenizer rate.

Encapsulation efficiency (EE) of drug-loaded nanodroplet emulsions was assessed using centrifugation at 4°C and 15 000 rpm for 40 minutes (Universal320R, Hettich). The purpose was to separate encapsulated drug from non-encapsulated drug. Subsequently, the supernatant, which contained the non-encapsulated drug, was carefully collected, ensuring minimal disturbance to the nanodroplet pellet. The quantification of non-encapsulated berberine chloride was achieved by measuring its absorbance at 418 nm with a UV-Vis spectrophotometer (UV-5100PC, Shanghai Metash Instruments Co., Ltd., China) and comparing the results to a standard curve. To calculate EE, the amount of non-encapsulated drug concentration was divided by the total amount of drug used for the nanodroplet preparation, as shown in the equation:

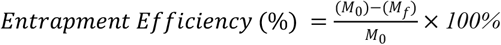

Here, M0 represents the total amount of drug added to the formulation, while Mf signifies the quantity of free drug in the supernatant, determined through a calibration curve constructed using standard solutions with known drug concentrations.

#### 2.3.5 Stability of Nanodroplets

Nanodroplets, suspended in phosphate-buffered saline (PBS) at pH 7.4, were incubated at 4°C for specific time intervals to evaluate their stability. We tracked changes in nanodroplet size and drug entrapment efficiency over a one-month period.

#### 2.3.6 In Vitro Evaluation of Drug Release

##### 2.3.6.1 Evaluation of Drug Release under Passive Conditions

The kinetics of drug release from nanodroplets were investigated using the dialysis method. For the experiment, 2 ml of optimized nanodroplet emulsion containing berberine chloride was placed in a dialysis bag (molecular weight cut-off 12000 Da), which was then immersed in 10 ml of PBS (pH 7.4) and citrate buffer (pH 5.5). Conditions were maintained at a constant temperature of 37° C and a stirring speed of 100 rpm using a magnetic stirrer. At specific time intervals (0–8 and 24 h), 2 ml of buffer was withdrawn and replaced with an equal volume of fresh buffer. The release of berberine chloride was quantified using a UV– vis spectrophotometer (UV-5100PC, Shanghai Metash Instruments Co., Ltd., China) at 418 nm. Berberine chloride release behavior was evaluated at two different pH values (7.4 and 5.5) to mimic physiological conditions and the slightly acidic extracellular matrix of most tumors. Tumor tissues often exhibit a lower pH compared to healthy tissues, typically ranging from 6.5 to 7.0, with some studies suggesting even lower values. The extracellular pH in the tumor microenvironment tends to be more acidic, while intracellular pH may vary, being closer to neutral or slightly alkaline in certain tumor cells. In order to determine the accumulated release of berberine chloride from nano droplets, the following formula was used:

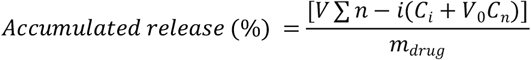

Where V is the sampling volume, V0 is the initial volume, Ci and Cn are the berberine chloride concentrations, i and n are the sampling times, and mdrug is the mass of berberine chloride in nanodroplets[15].

##### 2.3.6.2 Evaluation of Stimulated Drug Release

### 3.1 Effect of Polymer Concentration Variation

Alterations in particle size and drug loading were investigated by adjusting the polymer concentration within the emulsion system in formulations Aa, Ab, and Ac. A decrease in the gelatin weight percentage resulted in a progressive reduction in the particle size, which transitioned from 63 5.1 nm in formulation Aa to 458 nm in Ab and further to 281.7 nm in Ac, as shown in Table 2. Concurrently, a predictable decrease in drug loading was observed with decreasing polymer concentrations (2.3%, 2%, and 1.67% w/v), leading to a decrease in drug loading from 73.3% in Aa to 69.2% in Ab and finally to 66.8% in Ac. The polymer concentration within the emulsion system exerted a significant influence on both particle size and drug loading.

**Table 2.**
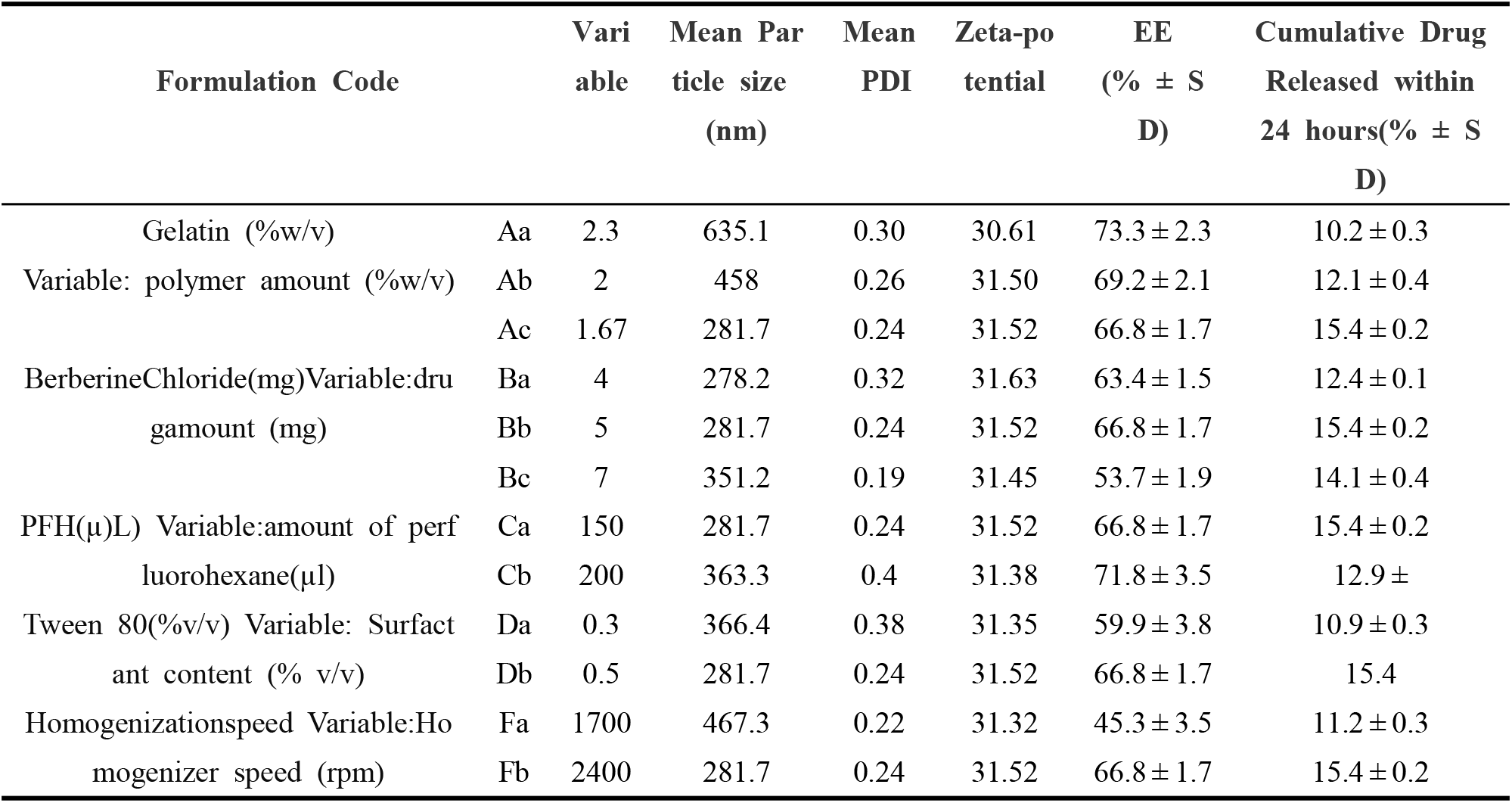
Effect of different formulations and structural parameters on the average particle size, polydispersity index (PDI), zeta potential, encapsulation efficiency (EE), and cumulative drug release. All components of the above formulations were added to 3 ml of deionized water.

| Formulation Code | | Variable | Mean Particle size<br>(nm) | Mean PDI | Zeta-potential | EE<br>(% ± S D) | Cumulative Drug<br>Released within<br>24 hours(% ± S D) |
| --- | --- | --- | --- | --- | --- | --- | --- |
| Gelatin (%w/v)<br>Variable: polymer amount (%w/v) | Aa | 2.3 | 635.1 | 0.30 | 30.61 | 73.3 ± 2.3 | 10.2 ± 0.3 |
| | Ab | 2 | 458 | 0.26 | 31.50 | 69.2 ± 2.1 | 12.1 ± 0.4 |
| | Ac | 1.67 | 281.7 | 0.24 | 31.52 | 66.8 ± 1.7 | 15.4 ± 0.2 |
| BerberineChloride(mg)<br>Variable:drug amount (mg) | Ba | 4 | 278.2 | 0.32 | 31.63 | 63.4 ± 1.5 | 12.4 ± 0.1 |
| | Bb | 5 | 281.7 | 0.24 | 31.52 | 66.8 ± 1.7 | 15.4 ± 0.2 |
| | Bc | 7 | 351.2 | 0.19 | 31.45 | 53.7 ± 1.9 | 14.1 ± 0.4 |
| PFH(μL) Variable:amount of perfluorohexane(μl) | Ca | 150 | 281.7 | 0.24 | 31.52 | 66.8 ± 1.7 | 15.4 ± 0.2 |
| | Cb | 200 | 363.3 | 0.4 | 31.38 | 71.8 ± 3.5 | 12.9 ± |
| Tween 80(%v/v) Variable: Surfactant content (% v/v) | Da | 0.3 | 366.4 | 0.38 | 31.35 | 59.9 ± 3.8 | 10.9 ± 0.3 |
| | Db | 0.5 | 281.7 | 0.24 | 31.52 | 66.8 ± 1.7 | 15.4 |
| Homogenizationspeed<br>Variable:Homogenizer speed (rpm) | Fa | 1700 | 467.3 | 0.22 | 31.32 | 45.3 ± 3.5 | 11.2 ± 0.3 |
| | Fb | 2400 | 281.7 | 0.24 | 31.52 | 66.8 ± 1.7 | 15.4 ± 0.2 |

A nanoemulsion of nanodroplets was first applied onto a culture dish serving as a scaffold to study the impact of ultrasound waves on the release of berberine chloride. The nanoemulsion was then exposed to low-frequency ultrasound (150 kHz, 0.2 W/cm^2, continuous mode) using a converter (SM3678B, Siemens, Shanghai, China) placed 0.5 cm away from the dish, equipped with probes having a cross-sectional area of 5 cm^2. Similarly, high-frequency ultrasound (1 MHz, 0.2 W/cm^2, continuous mode) was also applied using a converter (Enraf Nonius, Rotterdam, Th e Netherlands) under the same experimental conditions. Varied ultrasound exposure durations of 30 seconds, 5 minutes, and 10 minutes were employed. During the experiment s, the effective radiation area (ERA) was consistently measured at 30 mm^2, in alignment with the geometric cross-sectional area of the 5 cm^2 probes. Following the exposure, the nanodroplet solution underwent centrifugation at 1 5000 rpm, and the concentration of the released drug in the supernatant was quantified using established methods. Subsequently, the drug release data obtained from both lo w and high frequencies were compared.

### 2.4 Statistical Analysis

All experiments were conducted in triplicate. Statistical analyses were performed using two-way ANOVA and Tukey’ s multiple comparison tests. Significance was reported at P < 0.05.

## 3. Results and Discussion

The effects of different processes and structural parameters on the average particle size, zeta potential, entrapment efficiency, and cumulative amount of drug released from different formulations of nanodroplets are shown in Table 2.

Lower polymer concentrations facilitated the production of smaller particles but resulted in reduced drug loading. I n emulsion systems, the polymer serves as a stabilizer, aiding in the maintenance of drug particle dispersion. Higher polymer concentrations, such as those in formulation Aa, provide more available polymer chains for stabilizing particles, resulting in larger particle sizes. An increased polymer concentration enhances the capacity for drug encapsulation, thus preventing or delaying the saturation of drug encapsulation and improving drug loading and encapsulation efficiency[16]. The release of BBR chloride from formulations Aa, Ab, and Ac over 24 h was 10.2%, 12.1%, and 15.4%, respectively. Notably, a greater percentage of berberine chloride was released from formulation Ac (with smaller droplets) than from formulation Aa (with larger droplets), indicating a slower rate of drug release from Ac. Larger droplets, typically associated with slower drug release rates, can impede drug diffusion due to increased polymer concentration and viscosity, resulting in longer diffusion paths for drug molecules to traverse from the particle to the surrounding aqueous phase. Consequently, drug molecules may exhibit a greater propensity to be trapped within the particles, leading to increased drug retention or reduced drug release. Formulations with larger particle sizes, o wing to higher polymer concentrations, displayed elevated drug retention, as previously reported[17]. Based on the nanodroplet size, encapsulation efficiency (EE), and drug release profile, formulation Ac with 1.67% w/v gelatin was identified as the optimal polymer concentration. Additionally, the polydispersity index (PDI) for all the samples ranged between 0.2–0.3. PDI serves as a parameter for characterizing the particle size distribution, including that of nanodroplets, with values between 0.1 and 0.3 considered reasonable for various applications, including drug delivery systems[18]. Table 2 presents the positive zeta potential values of the nanodroplets. Gelatin is amphoteric, and its zet a potential is influenced by the pH of the surrounding medium. Under neutral conditions, gelatin tends to exhibit a near-neutral zeta potential. However, in practical drug deli very applications where the pH of the medium deviates from the isoelectric point (pI) of gelatin, gelatin nanodroplets typically display a slightly positive zeta potential in aqueous solutions. Zeta potential measurements within the range of 30–32 were consistent across different formulations, indicating minimal influence on zeta potential due to formulation alterations[19].

### 3.2 Effect of Drug Loading Variation

The particle sizes of the core-shell nanocarriers increased with increasing drug concentration. Specifically, the recorded sizes for drug loadings of 4 mg, 5 mg, and 7 mg were 278.2 nm, 281.7 nm, and 351.2 nm, respectively. This observed increase in particle size indicates that greater drug loading leads to the formation of larger nanocarrier structures. Interestingly, the entrapment efficiencies displayed an unexpected trend. At drug loading concentrations of 4 mg, 5 mg, and 7 mg, the entrapment efficiencies were 63. 4 ± 1.5%, 66.8 ± 1.7%, and 53.7 ± 1.9%, respectively. Contrary to the particle size results, the entrapment efficiency decreased with increasing drug concentration. This unexpected finding suggested that factors beyond the drug loading concentration may influence the encapsulation process. The increase in particle size with increasing drug loading can be attributed to the increased amount of drug incorporated into the nanocarrier system. However, the decrease in entrapment efficiency with increasing drug concentration is intriguing and warrants further investigation. It is conceivable that at higher drug concentrations, the drug molecules may have a propensity to aggregate and precipitate on the shell surface, thus reducing their incorporation into the core-shell nanocarriers. As expected, the formulation with a higher entrapment efficiency (Ba formulation) demonstrated greater cumulative drug release within the initial 2 4-hour period (refer to Fig. 1b). This finding suggested a correlation between entrapment efficiency and drug release kinetics in the studied system. This can be attributed to the improved retention of the drug within the nanocarrier, resulting in a more efficient drug release process[20].

**Figure 1.**
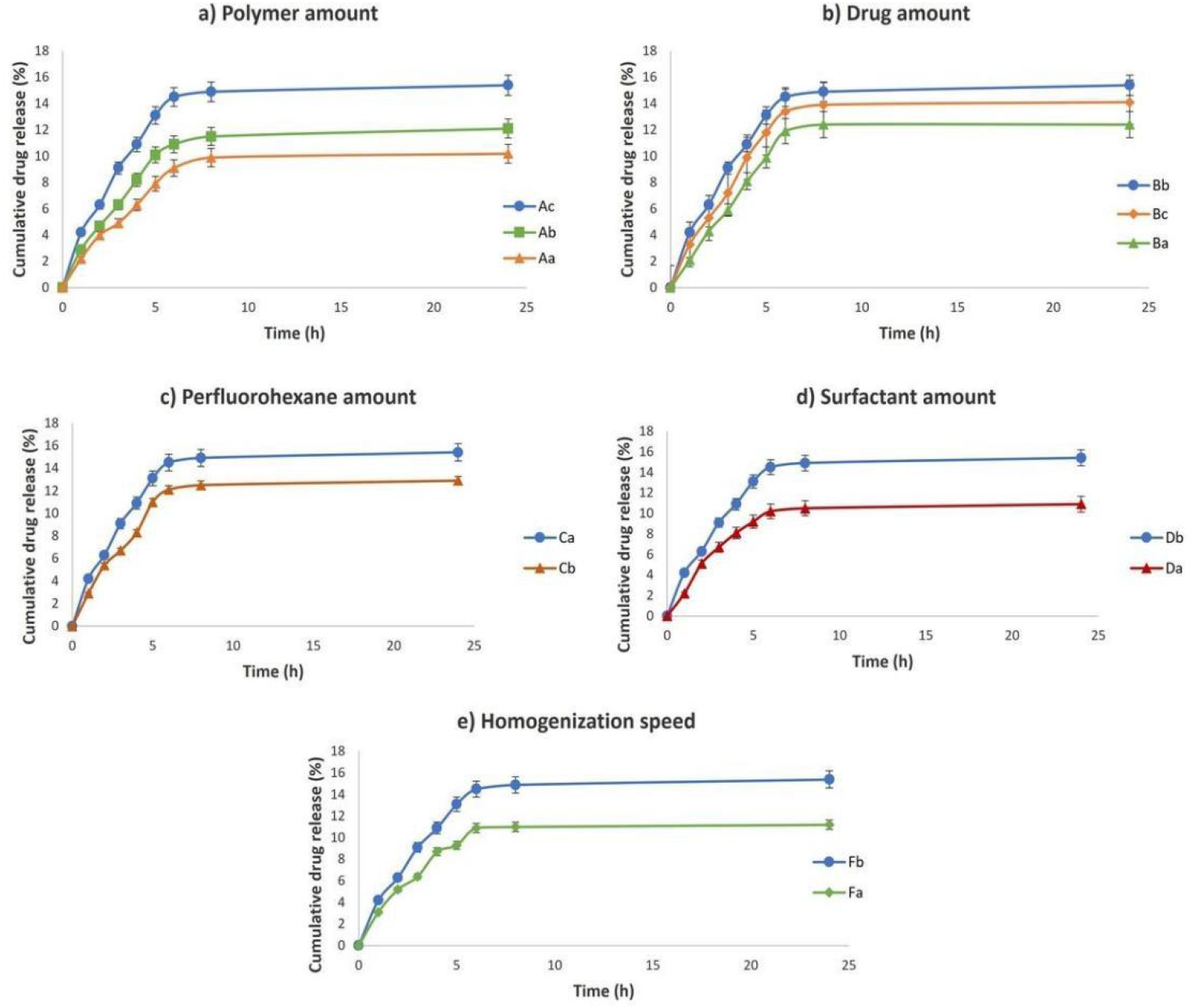
Cumulative drug released percentage within 24 hours in different formulations. Variables included a) polymer amount, b) drug amount, c) perfluorohexane amount, d) surfactant amount, and e) homogenization rate. Polymer amount, perfluorohexane amount, surfactant amount, and homogenization rate showed significant effects on drug-released profiles.

### 3.3 Effect of Perfluorohexane Variation

Perfluorohexane plays a pivotal role in core-shell nanocarriers as the core material, offering the ability to generate microbubbles upon ultrasound stimulation, thereby enhancing contrast and enabling targeted therapy. Due to its low surface tension, high volatility, biocompatibility, and chemical stability, perfluorohexane is indispensable for efficient dispersion, controlled release, and gas exchange, making it a versatile and invaluable component in nanocarrier design. This study investigated the impact of increasing the amount of perfluorohexane, a critical core component, on the size of nanodroplets and drug loading. The results revealed that increasing the perfluorohexane content led to an increase in the particle size, as shown in Table 2. Nanodroplets derived from the Ca and Cb formulations exhibited mean sizes of 281.7 nm and 363.3 nm, respectively. This surge in particle size can be ascribed to emulsions containing a higher oil phase content, namely, perfluorohexane. An elevated oil phase content may result in insufficient surfactant availability to adequately envelop the surface of each droplet and impede their fusion. Consequently, larger droplets are generated, culminating in a reduced specific surface area and increased particle size[21]. Surprisingly, variations in perfluorohexane content appeared to minimally impact drug loading. This suggests effective entrapment of the majority of drugs within the gelatin shell. This observation implies that the drug encapsulation process is governed primarily by the properties of the gelatin shell rather than by the quantity of perfluorohexane in the core[22].

With the increase in the oil phase, the drug release rate decreased from 15.4% to 12.9% in Ca and Cb, respectively (refer to Fig. 1c). A decrease in the specific surface are a resulting from an increase in nanocarrier size can influence drug release kinetics. A reduced surface area may impede drug diffusion and decelerate drug release from the nanodroplets, resulting in a diminished drug release rate[2 3]. These findings underscore the significance of considering the composition and formulation parameters of drug delivery systems. The quantity of perfluorohexane exerts a critical influence not only on particle size but also on drug release kinetics. Optimizing formulation parameters can facilitate the attainment of desired drug release profiles and enhance the therapeutic efficacy of the delivery system. Augmenting the perfluorohexane content results in nanodroplets with larger cores and thinner shells. Thinner shells and larger particle sizes lead to lower drug loading, while smaller nanodroplets necessitate less energy for microbubble conversion, consequently reducing heat generation following ultrasound stimulation[24].

### 3.4 Influence of Tween 80 Concentration

The choice and concentration of surfactant play a pivotal role in shaping the particle size of nanocarriers. Surfactant molecules serve as steric or electrostatic barriers, impeding the aggregation or coalescence of nanodroplets and resulting in smaller particle sizes. Moreover, surfactants can modulate emulsion droplet size, thereby impacting the final particle size of nanocarriers. Tween 80, also known as polysorbate 80, is a nonionic surfactant renowned for its exceptional emulsifying properties, rendering it highly effective in stabilizing oil-in-water emulsions. By lowering the interfacial tension between the oil and water phases, Tween 80 facilitates the formation of stable emulsions characterized by small droplet sizes[25]. This attribute proves particularly advantageous in nanocarrier preparation, facilitating the attainment of uniform and stable nanodroplet dispersions. Additionally, the lipophilic segment of Tween 80 enhances the solubility and dispersibility of poorly water-soluble drugs such as berberine chloride in aqueous media. Thro ugh micelle formation or drug solubilization, Tween 80 augments drug bioavailability and facilitates drug encapsulation within the nanocarrier system[26]. To probe the influence of varying surfactant concentrations on the particle size, drug loading rate, and drug release kinetics, two distinct concentrations of Tween 80 were used. Manipulating the surfactant concentration in the formulation allows for the modulation of the particle size and encapsulation efficiency. Increasing the surfactant concentration (0.3% and 0.5% volume of Tween 80) yielded smaller particle sizes in the Da (366.4 nm) and Db (281.7 nm) formulations, which was coupled with a heightened encapsulation efficiency in the Da (59.9%) and Db (66.8%) formulations. An increase in the percentage of Tween 80 promoted the formation of smaller and more uniform nanodroplets. By acting as a surfactant, Tween 80 diminishes the interfacial tension between the oil and water phases, facilitating superior dispersion and stabilization of hydrophobic compounds within the aqueous milieu. This phenomenon fosters the generation of diminutive droplets or nanodroplets during emulsification or self-assembly processes. Moreover, higher concentrations of Tween 80 enhanced the encapsulation efficiency of berberine chloride (refer to Table 2). Surfactant molecules engage with hydrophobic compounds, forming micelles or intricate structures that facilitate their entrapment within the nanocarrier system. This amplifies the loading capacity while mitigating potential drug loss during formulation[27]. The drug release profiles of the aforementioned formulations over 24 hours were 10.9% and 15.4%, respectively (see Fig. 1d). As expected, drug release was anticipated to be more pronounced in the Db formulation owing to the smaller size of the nanocarriers, as elucidated in Sectio n 3.3.

### 3.5 Impact of Homogenizer Speed

The homogenizer plays a pivotal role in the emulsification and dispersion of components to achieve a stable and uniform formulation. By applying mechanical shear forces, the homogenizer effectively breaks down the immiscible components, such as perfluorohexane (oil phase), and aqueous components (berberine chloride, Tween 80, gelatin). This process overcomes interfacial tension and generates smaller droplets of the oil phase dispersed in the aqueous phase. In our study, we explored two homogenization rates, 1 7000 rpm and 24000 rpm, to elucidate their effects on nanodroplet synthesis. It was observed that the particle size decreased with increasing velocity, aligning with expectations (refer to Table 2). The high-speed rotation of the homogenizer induces intense shear forces, fragmenting larger droplets into smaller droplets. Smaller droplets afford a larger interfacial area, facilitating the encapsulation of hydrophobic berberine chloride within perfluorohexane droplets and enhancing formulation stability. Moreover, increasing the homogenization speed to 24000 rpm corresponded to an increase in the encapsulation efficiency to 66.8%. A heightened homogenizer speed fosters improved emulsification, enhanced mixing, and optimized surface coverage, consequently bolstering drug encapsulation within nanodroplets[2 8]. Smaller droplets provide more opportunities for hydrophobic berberine chloride molecules to partition and become encapsulated within the oil phase (perfluorohexane) of nanodroplets, thereby enhancing encapsulation efficiency. Enhanced mixing ensures better distribution of the hydrophobic drug within the oil phase, augmenting drug encapsulation. Moreover, increased homogenizer speeds promote superior coverage of droplet surfaces by surfactant (Tween 80) molecules. This facilitates the effective adsorption and arrangement of surfactant molecules at the oil-water interface, culminating in a stable and well-structured interfacial layer. Improved surface coverage stabilizes nanodroplets, prevents coalescence, and enhances hydrophobic drug encapsulation[29]. Nanodroplets synthesized at 24000 rpm exhibited a greater release rate (15.4%) than larger nanodroplets obtained at 17000 rpm (11.2%) after 24 hours (refer to Fig. 4). This observation aligns with expectations, as smaller nanodroplets, as discussed in Section 3.3, are expected to exhibit greater drug release due to their smaller size.

### 3.6 Physicochemical Characterization of Optimal Formulation

Among the various formulations investigated for synthesizing nanodroplets encapsulating berberine chloride within a gelatinous shell, the Ac formulation emerged as the optimal candidate for further experimentation, with an average particle size of 281.7 nm and an encapsulation efficiency of 66.8%. The size and morphology of the drug-loaded nanodroplets were meticulously examined using both dynamic light scattering (DLS) and transmission electron microscopy (TEM) (Fig. 2b-c). TEM imaging revealed nanodroplets with a distinctive semicircular shape and an approximate size of 250 nm, which is consistent with the dimensions obtained from DLS measurements. Additionally, DLS analysis exhibited a favorable polydispersity index, indicating uniformity in size distribution. Notably, transmission electron microscopy (TEM) revealed the core-shell structure of the nanodroplets, with a discernible perfluorohexane (PH F) core enveloped by a gelatin shell. The presence of an optimal amount of perfluorohexane facilitated the formation of well-defined nanodroplets that were readily identifiable in the TEM micrograph[30]. Further insights into the structural integrity and chemical composition of the nanodroplets were obtained through Fourier transform infrared (FTIR) spectroscopy analysis. FTIR spectra of gelatin typically exhibit prominent amide bands, with the amide I band (1650-1630 cm−1) corresponding to C=O stretching vibrations of amide bonds in the protein backbone and the amide II band (1560-1530 cm−1) representing N-H bending and C-N stretching vibrations. Interestingly, the FTIR spectrum of the nanodroplet formulation displayed these characteristic gelatin peaks with minimal shifts, suggesting the preservation of the secondary structure of the gelatin in the presence of berberine chloride [31]. Distinct peaks attributable to berberine chloride were also evident in the FTIR spectrum, including the stretching vibrations of aliphatic carbon−hydrogen bonds (2920-2930 cm−1), aromatic C=C ring s (1537 cm−1), and C–H vibrations (1030 cm−1). Importantly, the presence of these characteristic peaks in the nano droplet spectrum confirmed the successful encapsulation of BBR chloride within the gelatin matrix without compromising its chemical integrity [32]. In summary, the physicochemical characterization of the optimal Ac formulation underscores its suitability as a promising nanocarrier system for the targeted delivery of berberine chloride, offering insights into its structural morphology, composition, and stability.

**Figure 2.**
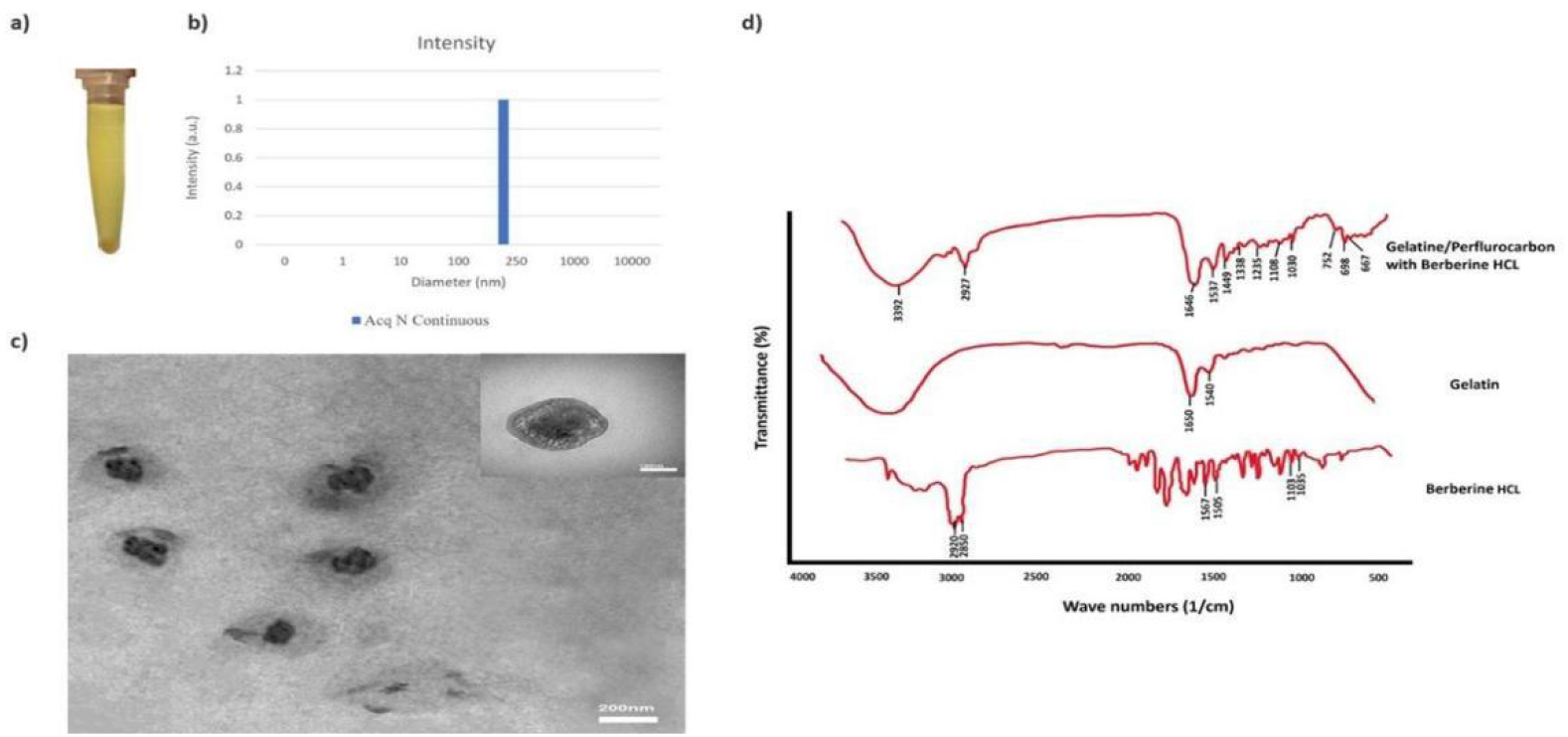
Physicochemical characterization of optimum formulation (Ac). a) Precipitated berberine chloride-loaded nanodroplets after centrifugation that separated for further analyses. b) the size distribution of nanodroplets obtained from DLS. The graph showed the particle size and high mono dispersity of the nanodroplets. c) TEM images of gelatin nanodroplets containing berberine chloride. Nanodroplets had a distinct core material (PHF) surrounded by a gelatin shell. High-magnitude detail inset: Inset in the top-left corner showcases a high-magnitude view of one nanodroplet within the TEM image. d) The FTIR spectrum of Ac formulation besides the drug and gelatin.

**Figure 3.**
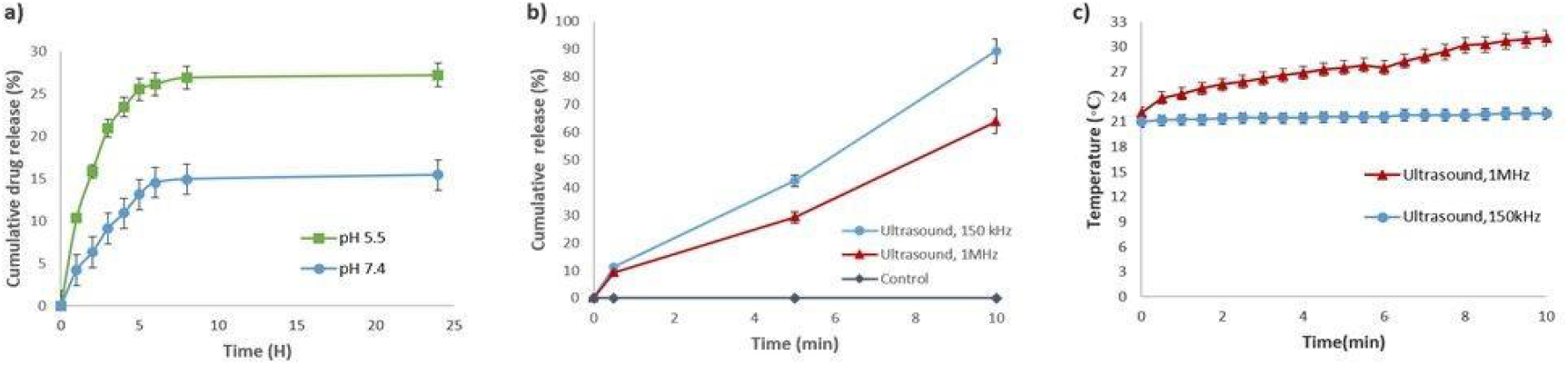
The comparison between passive and stimulated drug release of the optimum formulation and heat generation after ultrasound stimulation. a) Berberine chloride release profile from AC formulation at neutral pH (7.4) and acidic pH (5.5). The acidic condition changed the release profile b) The active release profile of berberine chloride from the optimal formulation (Ac) under low (150 k Hz) and high (1MHz) frequency ultrasound for 10 minutes. c) The temperature changing in the Ac formulation caused by the radiation of 1 MHz and 150 kHz ultrasound frequencies within 10 minutes, measured in degrees Celsius.

**Figure 4.**
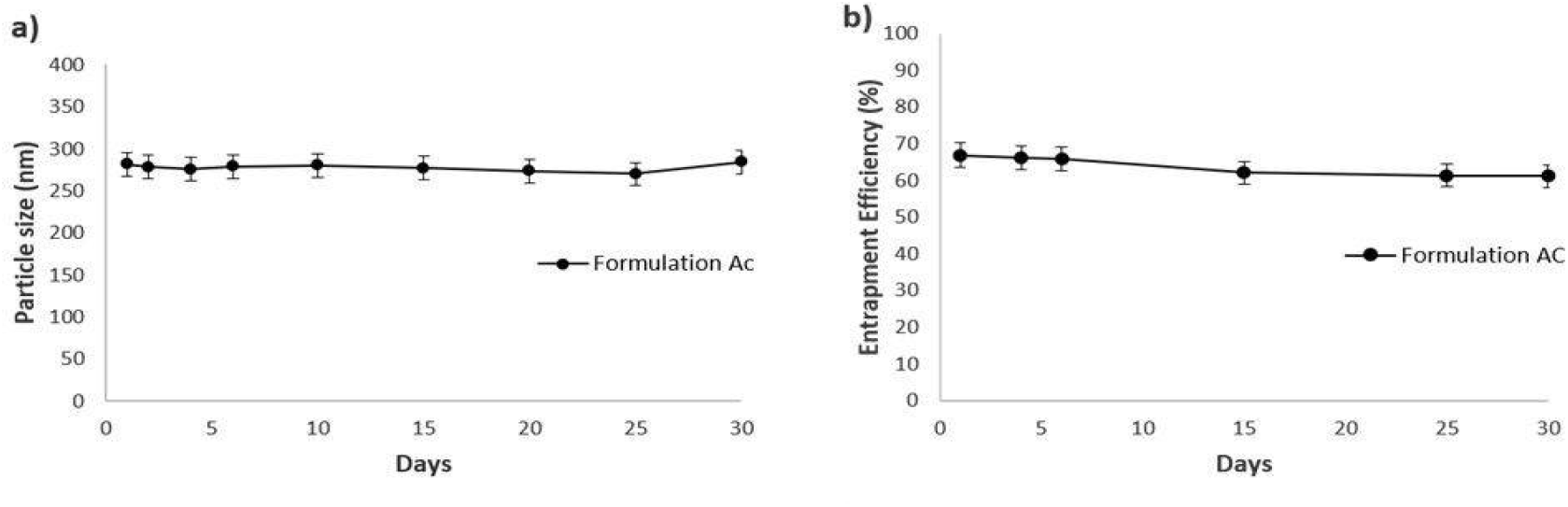
Changes in a) particle size and b) encapsulation efficiency of berberine chloride loaded gelatin system in AC formulation at a storage temperature of 4 ° C for predetermined time intervals. The trend showed stability of particle size and encapsulation efficiency over one month.

**Fig. 5.**
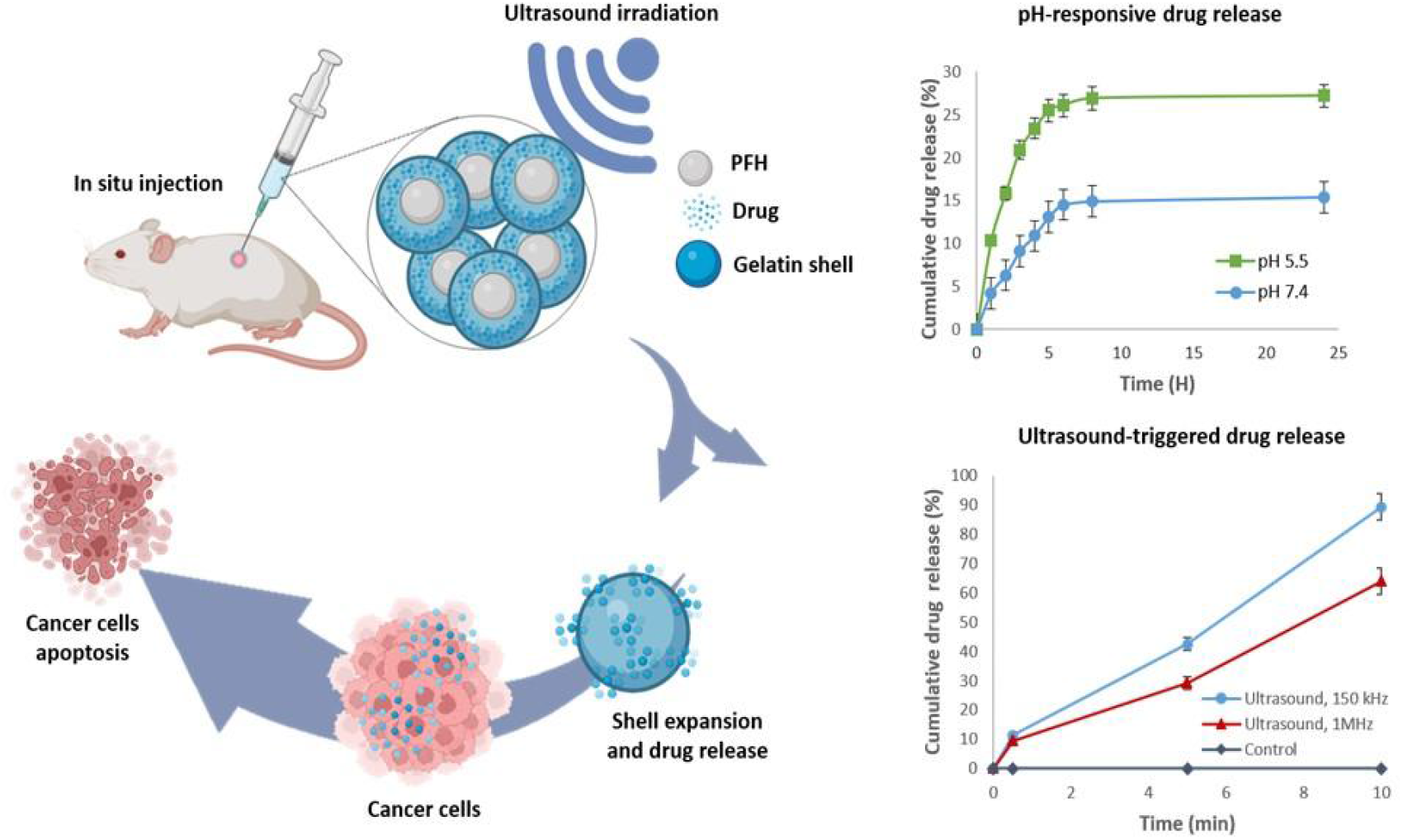
Schematic diagram of experimental process.

### 3.7 Passive Drug Release in Gelatin-Based Nanocarriers

Efficient drug delivery systems aim to strike a delicate balance between controlled drug release and minimizing passive diffusion, particularly in applications where precise temporal and spatial control of drug delivery is paramount. Achieving this balance is crucial for optimizing therapeutic outcomes while mitigating potential adverse effects associated with indiscriminate drug release. In the context of cancer therapy, minimizing passive drug release is of utmost importance to prevent off-target effects and reduce systemic toxicity.

In the Ac formulation utilizing gelatin as the shell material, passive drug release over 24 hours was observed to be 15.4% at neutral pH. Notably, this level of release was slightly higher compared to previous systems employing alginate-based shells. The observed differences may be attributed to the distinct structural properties of gelatin, which forms a denser and less porous matrix compared to alginate. This denser matrix potentially restricts the passive diffusion of drugs from the nanocarrier, leading to altered release kinetics [33].It is pertinent to acknowledge that the release kinetics of drugs from nanocarriers are influenced by various factors, including formulation composition, preparation method, and environmental conditions. Additionally, gelatin, as a protein-based material, may undergo conformational changes or degradation over time, impacting the permeability of the gelatin shell and subsequently influencing drug release dynamics.While gelatin-based nanocarriers may present limitations in passive release control, they offer distinct advantages in certain contexts. For instance, the Ac formulations exhibited significantly higher berberine chloride release at acidic pH (5.5, mimicking the endosomal/lysosomal environment) compared to neutral pH (7.4). This pH-responsive behavior can be attributed to the protonation of carboxyl groups in gelatin under acidic conditions, leading to alterations in the structure and permeability of the nanocarrier shell.In summary, the passive drug release profile of gelatin-based nanocarriers underscores the intricate interplay between formulation characteristics and environmental factors. While challenges in passive release control exist, the pH-responsive properties of gelatin-based nanocarriers hold promise for targeted drug delivery applications, particularly in acidic microenvironments associated with disease states such as cancer[34]. Continued research eff orts aimed at refining formulation strategies and elucidating underlying release mechanisms will further enhance the therapeutic potential of gelatin-based nanocarriers in precis ion medicine.

### 3.8 Ultrasound-Induced Drug Release

Upon initial injection into the body, perfluorocarbon nanodroplets exist in a liquid state. However, when exposed to ultrasound waves of specific frequency and intensity, these nanodroplets can undergo vaporization, transitioning into gas bubbles. This vaporization process is driven by the absorption of acoustic energy, causing rapid expansion and conversion into gas bubbles—a phenomenon known as acoustic droplet vaporization (ADV)[35].The mechanical forces generated during vaporization can disrupt or fragment the gelatin shell surrounding the nanodroplets, facilitating the release of encapsulated drugs or therapeutic agents. This process creates transient pores or openings in the gelatin shell, enabling controlled release and diffusion of the encapsulated drug, such as berberine chloride. Enhanced drug delivery via ultrasound-induced vaporization can potentially improve drug bioavailability, increase local drug concentration at the target site, and enhance therapeutic efficacy[36].

The frequency of ultrasound plays a crucial role in determining the vaporization threshold of perfluorocarbon nano droplets. Each type of nanodroplet has a specific vaporization threshold corresponding to particular ultrasound frequencies, typically ranging from tens of kilohertz (kHz) to megahertz (MHz). In our study, we investigated the ADV effect of the Ac formulation using thin needle injection and exposure to ultrasound at frequencies of 150 kHz and 1 MHz in a PBS solution at room temperature for 10 minutes. Our results demonstrated a significantly higher release of berberine chloride at 150 kHz (89.4%) compared to 1 MHz stimulation (63.9%) after 10 minutes[37].Lower ultrasound frequencies generally exhibit lower vaporization thresholds for ADV compared to higher frequencies, resulting in a higher percentage of nanodroplets undergoing vaporization. Additionally, lower frequencies promote stable cavitation, characterized by larger vaporized bubbles that undergo expansion and contraction cycles without immediate collapse. Stable cavitation can enhance drug release efficiency while minimizing potential bioeffects[38].It is important to consider safety aspects when utilizing ultrasound for therapeutic applications. Localized heat generation during ultrasound-induced vaporization can occur due to the conversion of acoustic energy into kinetic energy during bubble expansion and collapse. To ensure safety, it is recommended to limit the increase in local tissue temperature to no more than 2–4°C above baseline. Our findings indicated a clear temperature increase with 1 MHz radiation, highlighting the potential risk of tissue damage. Therefore, 150 k Hz radiation emerges as a safer and effective option for d rug release applications.

### 3.9 Stability of Nanodroplets

Figures 4a and b illustrate that over the course of one month at 4°C, there were negligible changes observed in both the size and the entrapment efficiency of the nanodroplets. This sustained stability holds promise for the advancement of an efficacious drug delivery platform. However, the acceptability of this one-month stability duration hinges significantly upon the specific requisites and intended application of the drug delivery system.

In a previous investigation11, alginate nanodroplets exhibit ed stability for up to four months. It is noteworthy that negatively charged nanoparticles like alginate often manifest heightened colloidal stability in aqueous milieu, as they repel each other by means of electrostatic repulsion. Conversely, proteins such as gelatin are generally more prone to degradation over time compared to polysaccharides, potentially resulting in diminished stability.In the current study, the selection of materials with a positive surface charge was prioritized to capitalize on enhanced cellular uptake and pH-responsive delivery. Positively charged nanoparticles are predisposed to more favorable interactions with negatively charged cell membranes. This electrostatic affinity ca n augment cellular uptake—a desirable trait when the objective is to ferry drugs or therapeutic agents into cellular interiors. Furthermore, gelatin nanodroplets exhibited pH-responsive behavior, affording the potential to refine drug delivery to specific tissues.

## 4 Conclusion

In this study, we have developed and optimized gelatin/perfluorohexane (PFH) nanodroplets loaded with berberine chloride for dual pH and ultrasound-triggered release. Through a systematic exploration of various formulation and process parameters, we identified an optimal formulation consisting of 1.67% w/v gelatin, 5 mg berberine chloride, 1 50 µL PFH, 0.5% v/v Tween 80, and 24,000 rpm homogenization speed. This formulation yielded nanodroplets with desirable characteristics, including a size of 281.7 nm, a polydispersity index of 0.24, and an encapsulation efficiency of 66.8%. Physicochemical characterization using TEM and FTIR confirmed the core-shell structure of the nanodroplets and successful encapsulation of berberine chloride.Our in vitro release studies demonstrated the pH-responsive behavior of the nanodroplets, with higher cumulative release under acidic conditions compared to neutral pH, attributed to the protonation of the gelatin shell. Furthermore, ultrasound stimulation induced rapid acoustic droplet vaporization and burst release of the encapsulated drug, particularly at a lower frequency of 150 kHz, which generated minimal heat. This finding suggests the potential of utilizing ultrasound for controlled drug release with reduced side effects. Overall, our findings highlight the promise of gelatin/PFH nanodroplets as a dual-triggered nanocarrier system for targeted drug delivery. By leveraging the synergistic effects of pH responsiveness and ultrasound stimulation, these nanodroplets offer enhanced drug release at specific sites while minimizing systemic side effects. Further studies, including in vitro cytotoxicity and in vivo evaluations, are warranted to validate the efficacy and safety of these nanocarriers for clinical applications. This study contributes valuable insights into the development of smart multifunctional nanocarriers for precision medicine.

## Supporting information

Highlights

## Acknowledgements

This work was supported by a project grant from National Natural Science Foundation of Chin a General Project (32360113;72274193;81903791); Innovation Training Program of the Chinese Academy of Sciences (20234001153;20234002921); Gansu Province National Innovation Training Program for College Students (20230105 8) and National Special Fund for the Construction of Industrial Technology System of Chinese Herbal Medicine (C ARS-21).

