## Supplementary material for "PH-SENSITIVE NANODROPLETS FOR CONTROLLED DELIVERY OF BERBERINE CHLORIDE": Highlights

Here are the highlights for my article in English, suitable for submission to Biochemical and Biophysical Research Communications:

1. Development of pH-Sensitive Nanodroplets: We have successfully developed pH-sensitive nanodroplets based on gelatin/perfluorohexane (PFH) that enable precise control over the release of berberine chloride.

2. Optimization of Formulation Parameters: Through meticulous optimization of preparation parameters, we achieved nanodroplets with a mean particle size of 281.7 nm and a drug encapsulation efficiency of 66.8 ± 1.7%, demonstrating excellent preparative characteristics.

3. Enhanced Drug Release with Ultrasound Stimulation: Our study reveals a significant enhancement in drug release with ultrasound stimulation, particularly at a frequency of 150 kHz, providing strong support for the responsiveness of nanodroplets in drug delivery.

4. Long-Term Stability at Low Temperatures: Stability studies demonstrate that the nanodroplets maintain prolonged stability over one month at 4° C, underscoring their potential for practical applications in pharmaceutical formulations.

5. Integration of Traditional Medicine with Modern Drug Delivery: These pH-sensitive nanodroplets offer a promising platform for the controlled delivery of berberine chloride, facilitating the convergence of traditional Chinese medicine with advanced drug delivery strategies.

These highlights succinctly encapsulate the significant findings and contributions of our research, emphasizing its relevance and potential impact to the readership of Biochemical and Biophysical Research Communications.
